# Variation in cephalic neuromasts surface and cave-dwelling fishes of the family Amblyopsidae (Teleostei: Percopsiformes)

**DOI:** 10.1101/701490

**Authors:** Daphne Soares, Matthew L. Niemiller

## Abstract

Cave adaptation has led to unique sensory specializations to compensate for the lack of visual cues in aphotic subterranean habitats. As the role of vision is reduced or disappears, other sensory modalities become hypertrophied allowing cave-adapted organisms to successfully detect and interact their surrounding environment. The array of aquatic subterranean habitats, from fast-flowing streams and waterfalls, to quiet phreatic pools, presents a diverse palette to examine what possible sensory solutions have evolved against a backdrop of complete darkness. Mechanosensation is enhanced in many subterranean animals to such an extent that a longer appendage is recognized as a prominent troglomorphic adaptation in many metazoans. Fishes, however, not only interact with the environment using their fins, but also with specialized sensory organs to detect hydrodynamic events. We hypothesize that subterranean adaptation drives the hypertrophy of the mechanosensory lateral line, but that other environmental forces dictate the specific neuromast phenotype. To this end, we studied differences in the cephalic lateral line of the fishes in the North American family Amblyopsidae, which includes surface, cave-facultative, and cave-obligate species. None of the taxa we examined possessed canal neuromasts on the head. Primarily surface-dwelling species, *Chologaster cornuta* and *Forbesichthys agassizii*, possessed receded neuromasts throughout most of the head, with a few on papillae located in front of the nostrils and on ventral grooves on each side of the mouth. The cavefishes *Amyblopsis spelaea* and *Typhlichthys subterraneous* possessed papillate superficial neuromasts all over the head. We speculate that the change from the surface to the cave environment has led to papillate neuromasts in this group, which are likely shaped to detect the hydrodynamic characteristics of the boundary layer created by the swimming fish. Moving sensory organs from the surface of the body out into the boundary layer could increase sensitivity to high frequency stimuli created by prey, predators, and conspecifics.

## Introduction

Water currents are a pervasive feature of all aquatic environments and provide obvious cues influencing fish behaviors. Fishes themselves can also create water displacement and pressure fluctuations (Kalmijn 1989). Hydrodynamic stimuli provide important physical and biological information about the surrounding environment, and fishes have evolved a unique mechanosensory organ to detect hydrodynamic stimuli: the mechanosensory lateral line system (Bleckmann 1994; Bleckmann and Zelick 2009). The lateral line system is present in all fishes but varies in morphology and distribution.

The functional unit of the lateral line is the neuromast, which can be free standing on the skin (i.e., surface or superficial neuromasts) or in canals that are open to the environments via pores (i.e., canal neuromasts). The distribution of neuromasts across the body and head determines the ability of a fish to detect moving as well as stationary stimuli. Each object, be it a rock or a conspecific, creates a unique hydrodynamic signature. The information gained from these hydrodynamic images influences many aspects of fish behavior. For example, fish use their lateral line to examine novel objects (Teyke 1990; Deperera 2004), to detect prey (Hoekstra and Janssen 1985; Janssen 1999; Yoshizawa et al. 2010), to monitor the movement of conspecifics (Partridge et al., 1980; Faucher et al., 2010) and predators (McHenry et al., 2009), and to maintain position in flowing water (Sutterlin and Waddy 1975; Montgomery et al. 1997). Superficial and canal neuromasts differ in function and performance, but are overlapping in hydrodynamic selectivity. Physiological studies have shown that canal neuromasts respond to water movements that are produced by moving objects, such as prey, that is maximized in the direction of the canal axis (Denton and Gray 1989; Montgomery et al. 1994). Canal neuromasts typically occur in a distinct line at the base of a canal that runs and extends over the head and flanks. In contrast, superficial neuromasts are located on the surface of the skin and preferentially respond to the velocity of water flow that is not orthogonal to their orientation axis (Coombs and Montgomery 1994; Montgomery et al. 1994; van Netten and Kroese 1989) and appear to be best suited to encode flow velocity (Baker and Montgomery, 1999; Van Trump and McHenry 2013).

The principle sensory cell of the neuromast is the hair cell (Carton and Montgomery, 2004). These hair cells have a bundle of stereovilli that grow longer from one side of the apical surface to the other. A single true kinocilium occurs in the center on of the stereovilli bundle. The stereovilli of superficial hair cells are embedded in a gelatinous matrix called the cupula, and the cupula is drag coupled to the surrounding water. The motion of the hair cell bundles in the neuromast generates electrical potentials that are transduced into action potentials in afferent neurons (Mogdens and Bleckmann 2012). The relationship between cupula displacement and the amplitude of the receptor potential is linear and can reach saturation (Curcic-Blake and Van Netten 2006). These signals provide the central nervous system with information about water flow around the body.

The evolution of the lateral line system can be crucial for fishes to adapt to novel environments. For example, several studies have noted that species living in quieter, low-flow habitats tend to have more superficial neuromasts than species living in higher-flow habitats (Teyke 1990; Coombs et al. 1988; Janssen 2004). A similar response has been documented in some cavefishes, where blind cave-dwelling taxa have more superficial neuromasts than in related surface-dwelling populations that retain a functional visual system (Poulson 1963; Niemiller and Poulson 2010; Yoshizawa et al. 2010, 2014). It is possible that rather than being related to flow regime, morphological characteristics of the lateral line system may be an adaptive response to living in perpetual darkness in subterranean habitats. In Mexican blind cavefish, *Astyanax mexicanus*, superficial neuromasts are longer and more sensitive than those of related surface fish (Teyke 1990, Yoshizawa et al. 2014), which likely contributes to their ability to better detect flow. Although it is recognized that cavefishes generally exhibit hypertrophy of the lateral line system with respect to number and size of superficial neuromasts (Soares and Niemiller 2013; Niemiller and Soares 2015), studies on neuromast morphology of cavefishes are surprisingly few (Poulson 1963; Dezfuli et al. 2009; Jiang et al. 2016), with the most intensive work on *Astyanax* (Teyke 1990; Baker and Montgomery 1999; Montgomery et al. 2001; Yoshizawa et al. 2010, 2014).

We are interested in cephalic neuromasts specifically because they are crucial for the finetuning locomotion necessary for prey catching (Müller and Schwartz 1982, Tittel et al. 1984; Hoekstra and Janssen 1985, Janssen 1990; Janssen et al. 1995; Janssen 1996; Janssen and Corcoran 1998, Schwarz et al. 2011), and they may be organized in a conformation that leads to a mechanosensory fovea (Hoekstra and Janssen 1986, Coughlin and Strickler 1990). These neuromasts are very diverse in morphology and organization (Coombs et al. 1994; Beckmann et al. 2010), which are often related to the environment of the fish (Higgs and Furiman 1998; Ahnelt et al. 2004; Schmitz et al. 2014). Neuromast phenotypes can consequently be used as a diagnostic character for species delimitation (Kurawaka 1976, Reno 1966; Nelson 1972; Gosline 1974; Parin and Astakhov 1982; Stephens 1985; Webb 1989b; Arai and Kato 2003; Begman 2004). Functionally, cephalic neuromasts also contribute to rheotaxis (Baker and Montgomery 1999, Janssen, 2004) and schooling (Blaxter et al. 1983; Janssen et al. 1995; Pitcher 2001; Diaz et al. 2003),

Here, we examine and compare the morphology of cephalic neuromasts in a family of fishes that span the ecological gradient from surface to obligate cave inhabitation—the Amblyopsidae (Actinopterygii: Teleostei: Percopsiformes). The Amblyopsidae is comprised of six genera and nine species in eastern North America (Niemiller and Poulson, 2010; Chakrabarty et al. 2014; Armbruster et al., 2016). Although subterranean adaptation in fishes is quite common (Soares and Niemiller 2013; Niemiller and Soares 2015), this family is unusual in that most species in the family are stygobiotic (obligate subterranean). The swampfish (*Chologaster cornuta*) is a small, pigmented species that lives in swamps of the Atlantic Coastal Plain and is the only species in the family only found in surface habitats. All other recognized species are associated with karst terrain of the Interior Low or Ozark plateaus and are at least partially cave-adapted. The spring cavefishes, *Forbesichthys agassizii* and *F. papilliferus*, are facultative cave inhabitants generally occurring in spring-fed streams and also caves on occasion in the Interior Low Plateau karst region (Niemiller and Poulson 2010). All of other species in the family are obligate inhabitants of caves and have evolved a suite of morphological, physiological, and behavioral characters associated with subterranean life, most notably degeneration of eyes and reduction of pigmentation (Niemiller and Poulson 2010; Soares and Niemiller 2013): Northern Cavefish (*Amblyopsis spelaea*), Hoosier Cavefish (*A. hoosieri*), Alabama Cavefish (*Speoplatyrhinus poulsoni*), Ozark Cavefish (*Troglichthys rosae*), Eigenmann’s Cavefish (*Typhlichthys eigenmanni*), and Southern Cavefish (*Ty. subterraneus*). In this study, we describe the morphology of cephalic neuromasts using scanning electron microscopy in four species of amblyosids and compare among cave and surface taxa: *C. cornuta*, *F. agassizii*, *A. spelaea*, and *T. subterraneus*. We hypothesize that subterranean adaptation has lead to the hypertrophy of the cephalic mechanosensory lateral line in amblyopsid cavefishes.

## Materials and Methods

Individuals of four amblyopsid species were collected under scientific permits issued by the states of Tennessee (no. 1605) and Kentucky (no. SC1211135), USA in April 2012. We collected three individuals of *Forbesichthys agassizii* from a quiet pool (10 m^2^, mean depth 0.6 m, mud/silt substrate with abundant vegetation) of a spring run fed by Jarrell’s Spring, Coffee Co., Tennessee, USA; three individuals of *Chologaster cornuta* from Colly Creek, Bladen Co., NC; three individuals for each of the two cave-dwelling species: *Amblyopsis spelaea* from several quiet pools (20–150 m^2^, mean depth 1 m, silt/sand/cobble substrate) in Under the Road Cave, Breckinridge Co., Kentucky, USA and *Typhlichthys subterraneus* from several pools with some current (4–12 m^2^, mean depth 0.5 m, cobble/bedrock substrate) in L&N Railroad Cave, Barren Co., Kentucky, USA.

Four fishes we used for histology. Fish were euthanized by prolonged immersion in tricaine methane sulfonate (MS222, 300 mg/l pH 7.2) then perfused with 0.9% NaCl followed by Alcohol-Formalin-Acetic acid (AFA). Fish were decapitated and heads were placed in AFA overnight. Heads were decalcified (RDO Gold, 20:1 ratio) under slow agitation until soft (1-2 days), were then imbedded in paraffin (Paraplast X-tra), and cut at 10um thickness in a rotating microtome. Slides were rehydrated in a series of alcohol concentrations, stained with Cresyl violet for 5 minutes, rehydrated, cleared (Histoclear) and cover slipped with permount. Scanning electron microscopy (SEM) was used to differentiate and quantify the morphological features of superficial neuromasts among amblyopsid cavefishes. Fish were euthanized by prolonged immersion in tricaine methane sulfonate (MS222, 300 mg/l pH 7.2). Whole specimens were fixed in 2.5% glutaraldehyde, 2% parafomaldehyde in 0.1M phosphate buffer. Fish were dehydrated in a series of increasingly concentrated ethanol baths before being critically point dried and spur coated with 5 nm of gold-palladium in a Denton Vacuum Desk II. SEM samples were imaged in an AMRAY 1620D (Bedford, MA, USA) scanning electron microscope at 10 kV–30kV acceleration voltage. Scanning electron microscopy was performed at the Laboratory for Biological Ultrastructure at the University of Maryland. We defined the area of the neuromast as the surface that contains the entire footprint of the cupula and the sensory plate area, as the smaller region that contains only sterocilia. Measurements were done with ImageJ (open source, https://imagej.net). We sampled the entire head and operculum. We tested for differences in neuromast surface area, kinocilium length, and sensory plate neuromast area among species with analysis of variance followed by ad hoc Tukey HSD test using Vassarstats.net. Values reported are mean ± 1 standard deviation.

## Results

The heads of *Forbesichthys agassizii* (Figure 1) and *Chologaster cornuta* (Figure 2,7) were covered with stich-like structures, positioned dorsal-ventrally. The neuromasts on the dorsal and lateral part of the head were receded into pits. The ventral side of the jaw and rostrum in both of these species were populated with superficial neuromasts on papilla. The mandibular papillae were situated in grooves that run from the tip of the lower jaw to the edge of the mouth. The mean surface area of each superficial neuromast in *C. cornuta* was 272.9±45.0 μm_2_. The sensory plate, positioned in the center of the neuromast, was made up of about hundreds of hair cells that occupy 131.7 ±9.1 μm_2_, or 48.3% of the neuromast area. The polarity, from shorter to longer, of the stereocilia was perpendicular to the long axis of the neuromast. The length of the kinocilium was 3.5±1.5 μm. In *F. agassizii*, mean surface area was 320±40 μm^2^, with hair cells occupying 121.7±22.9 μm_2_, or 38% of the area of the sensory plate. The length of the kinocilium was 3.0± 1.5 μm. Each organ was raised above the skin on a fleshy papilla approximately 0.5 mm high. All cupulae were removed during processing.

**Figure 1.**
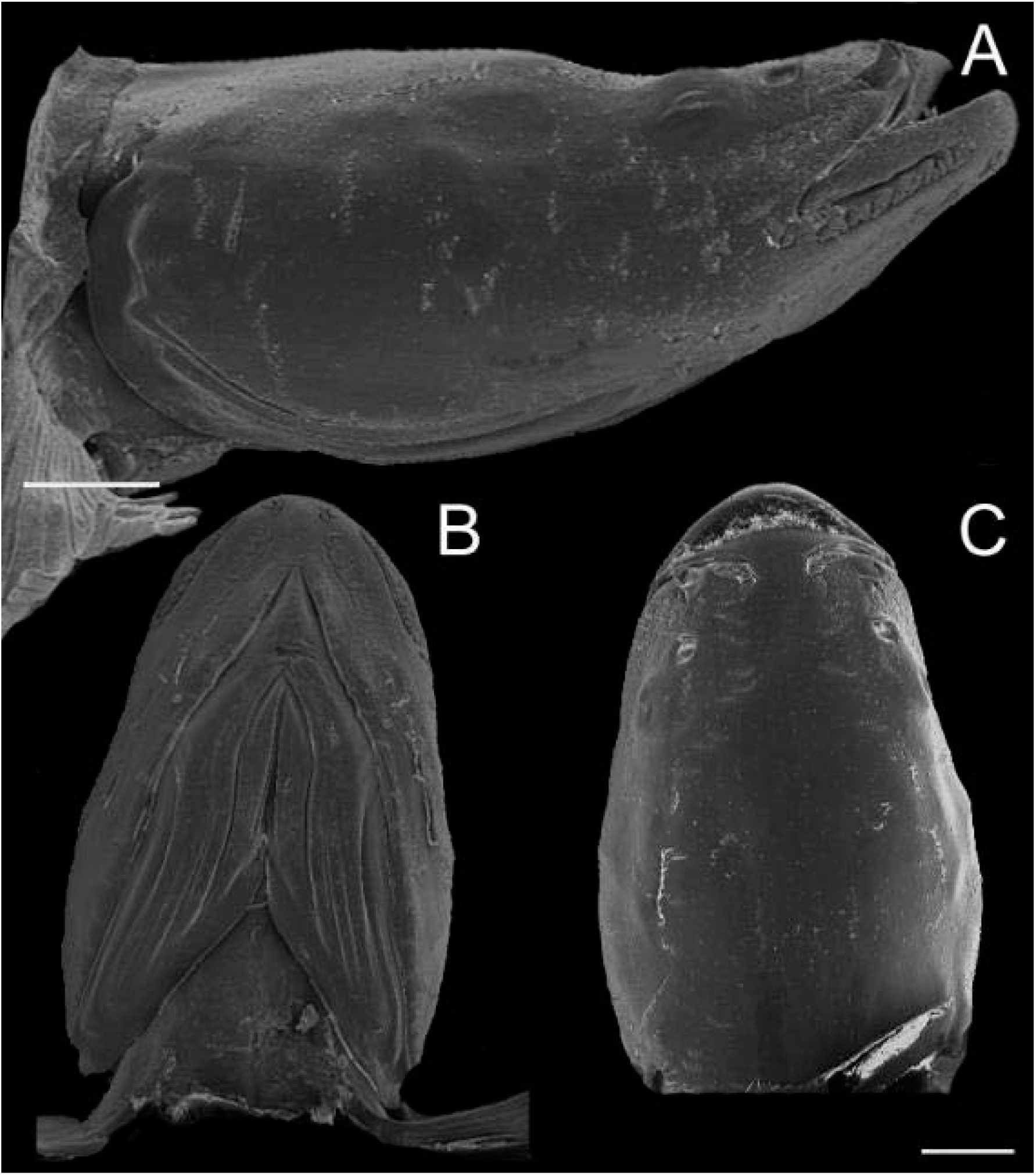
Lateral (A), ventral (B), and dorsal (C) SEM images of the head of *Forbesichthys agassizii*. Scale bar 1 mm.

**Figure 2.**
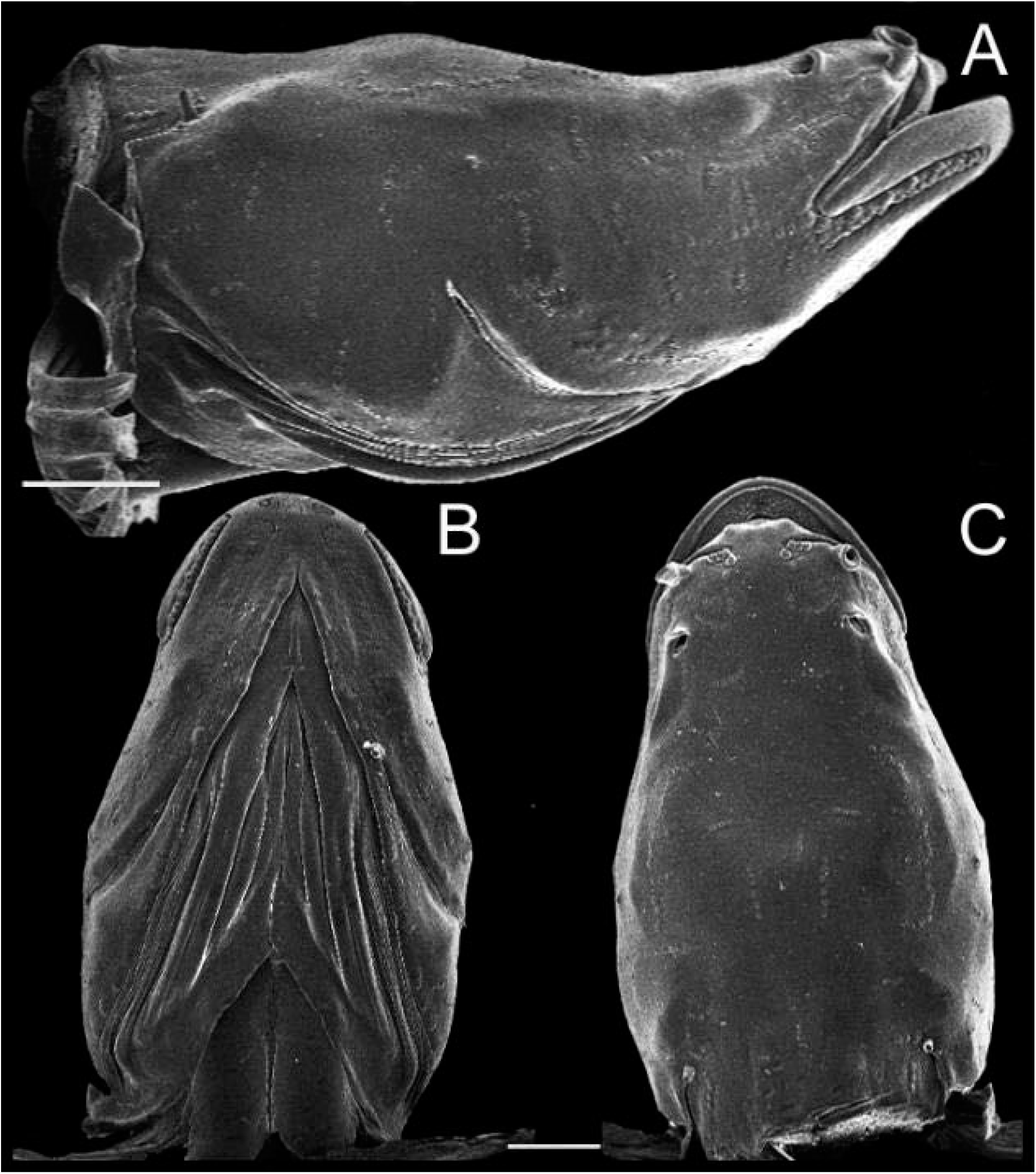
Lateral (A), ventral (B), and dorsal (C) SEM images of the head of *Chologaster cornuta*. Scale bar 1 mm.

**Figure 3.**
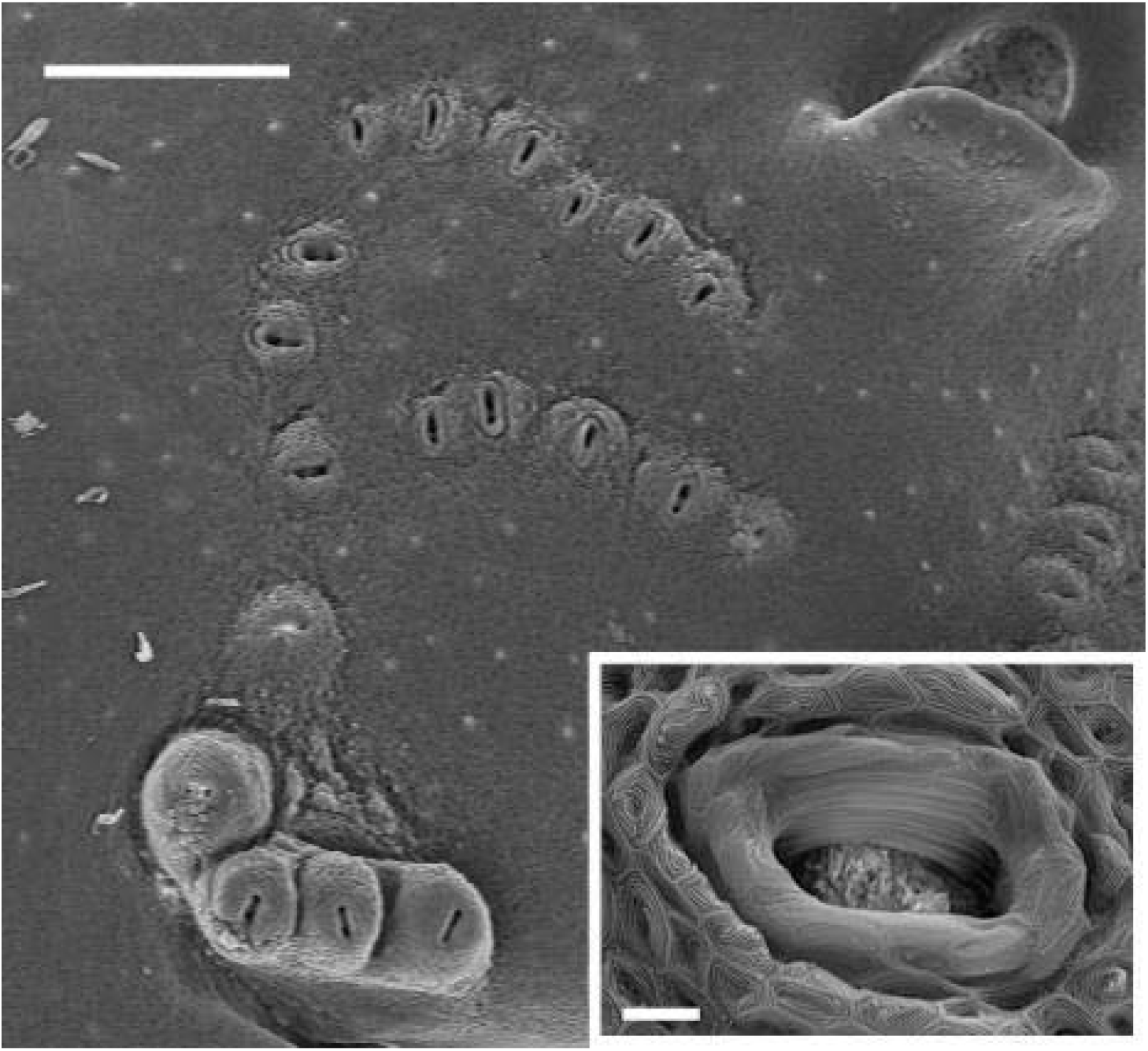
Receded neuromasts and neuromasts on papillae on the rostral end of the head of *C. cornuta*. Scale bar 250μm. Insert: stereocilia after the cupula was removed. Scale bar 10um.

Both species of cavefishes examined (*Typhlichthys subterraneus* and *Amblyopsis spelaea*) had more stiches than the two surface fishes (surface fishes= 18±3 vs cavefish= 25±6; Figure 4 and 5). The stiches were also longer in the dorsal-ventral dimension in *T. subterraneus*. All neuromasts in cavefishes had hair cells on papillae (height of papillla: *A. spelaea* = 0.55±0.006 mm; *T. subterraneus* 0.73±0.16 mm; Figure 6,7). In *A. spelaea*, neuromasts had a surface area of 759.1±133 μm^2^. The sensory plate of the neuromast had a mean area of 210.8±27.0 μm^2^, or 27% of the surface area of the neuromast. The kinocilium was 7.0±1.1 μm, and we were able to measure the cupula at 30.0± 6μm. In *T. subterraneus*, mean neuromast surface area resembled surface fishes at 224±59.7 μm^2^. The sensory plate area was 56.8±18.0 μm^2^, occupying 25% of the surface area of the neuromast. The length of the kinocilium was 5.4±0.9 μm.

**Figure 4.**
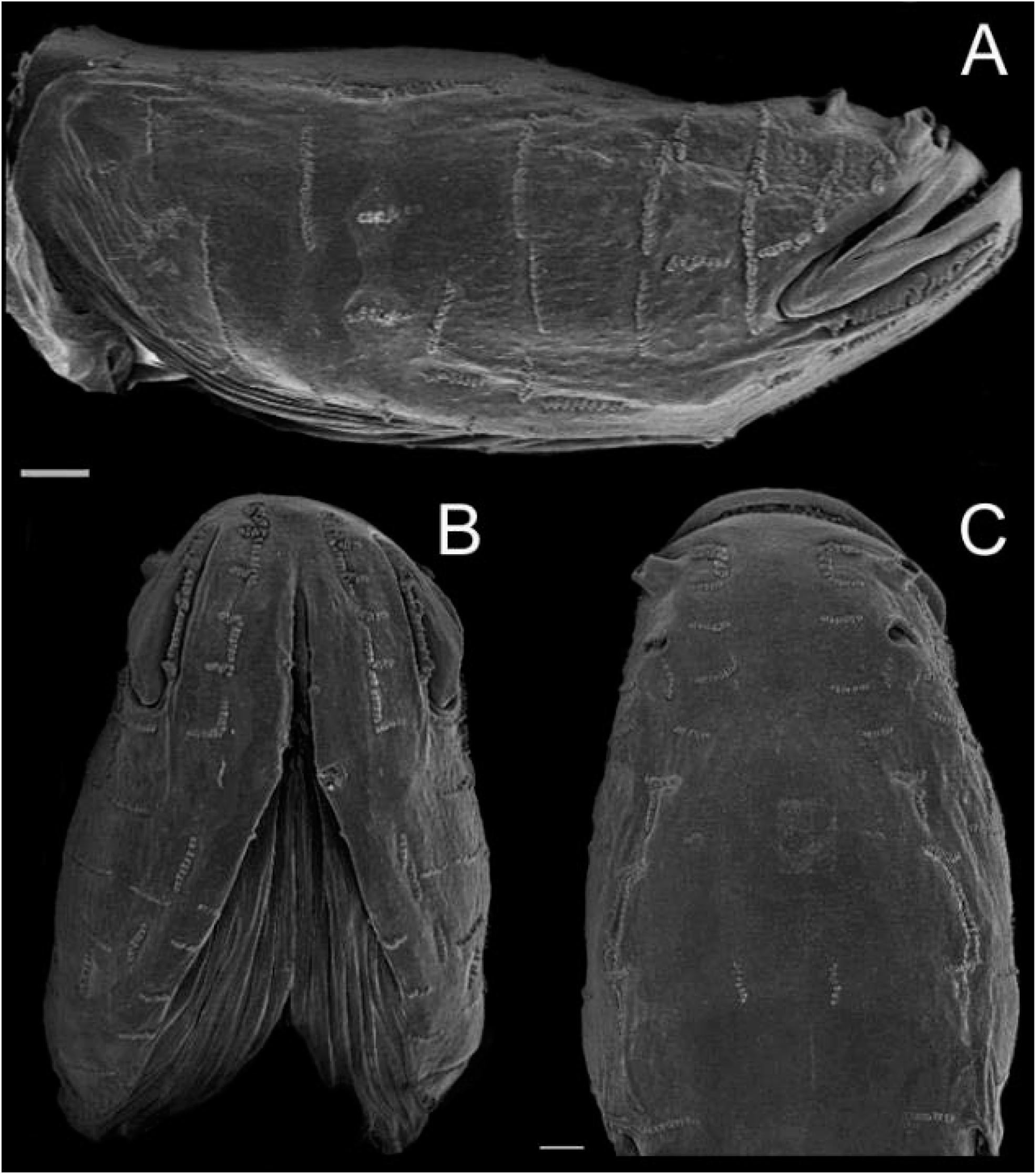
Lateral (A), ventral (B), and dorsal (C) SEM images of the head of *Typhlichthys subterraneus*. Scale bar 1 mm.

**Figure 5.**
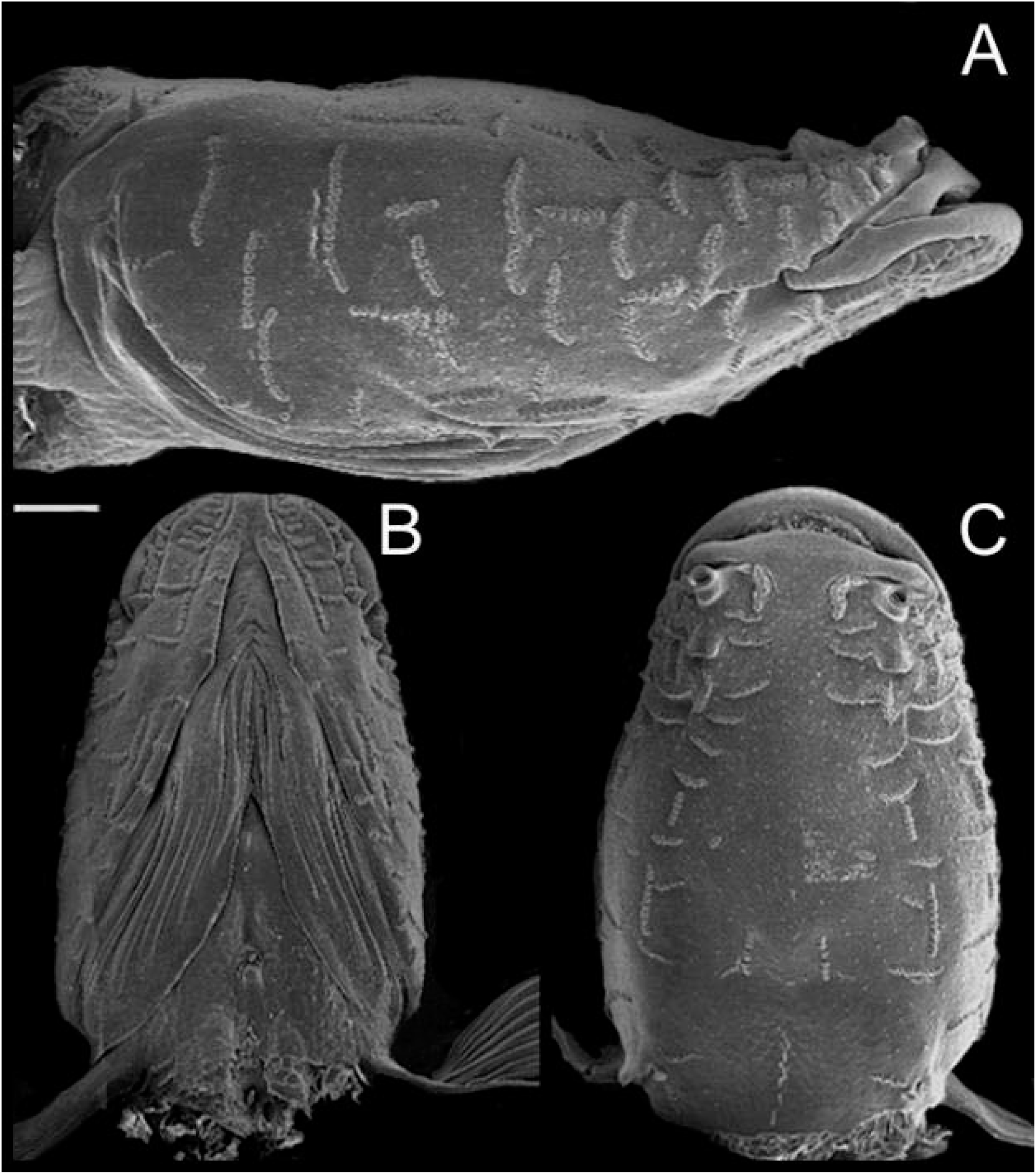
Lateral (A), ventral (B), and dorsal (C) SEM images of the head of *Amblyopsis spelaea*. Scale bar 1 mm.

**Figure 6.**
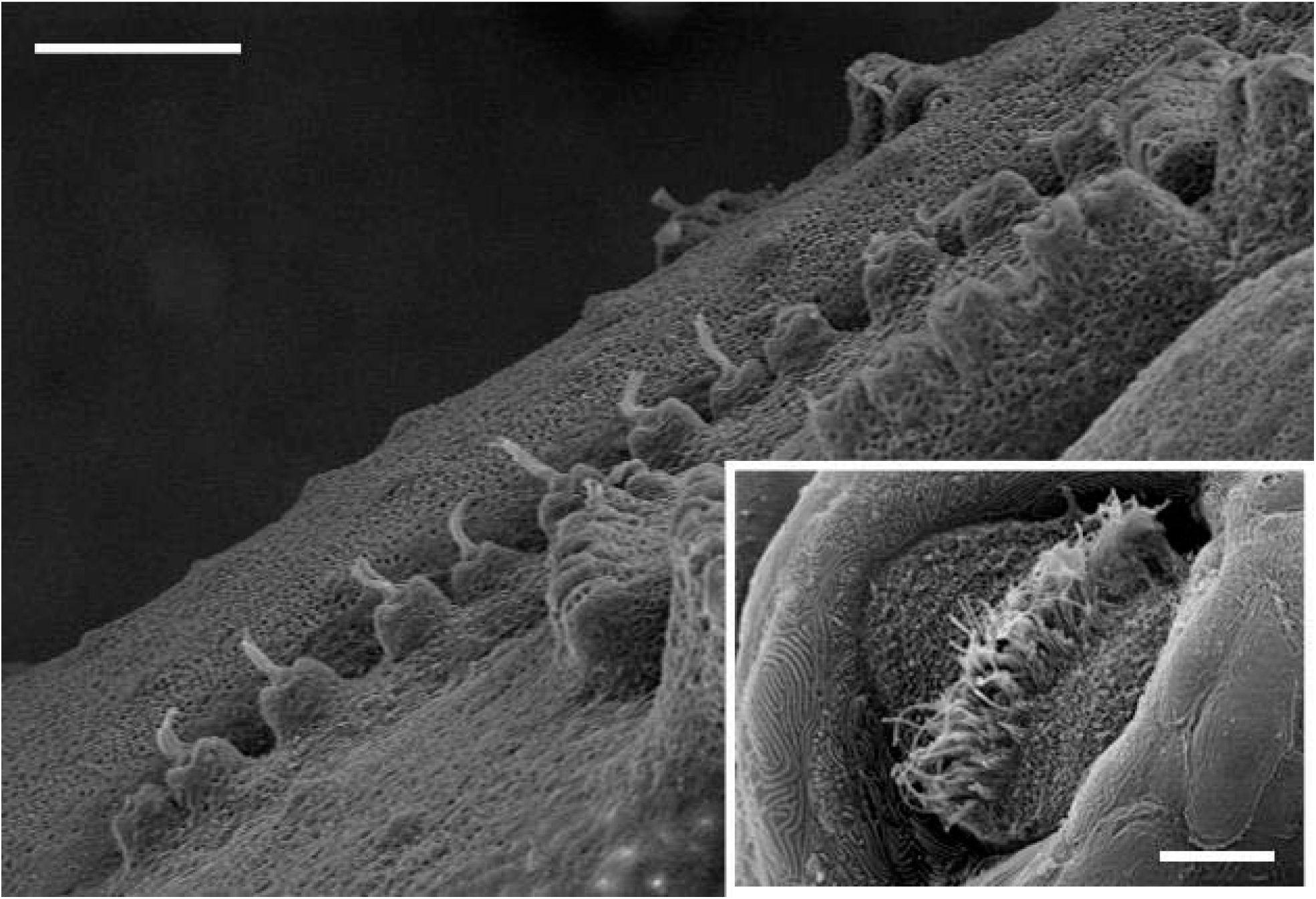
Neuromasts on papillae of *Amblyopsis spelaea* showing cupulae. Scale bar 250μm. Insert: stereocilia after the cupula was removed. The kinocilium is the longest of the bundle. Scale bar 10um.

**Figure 7.**
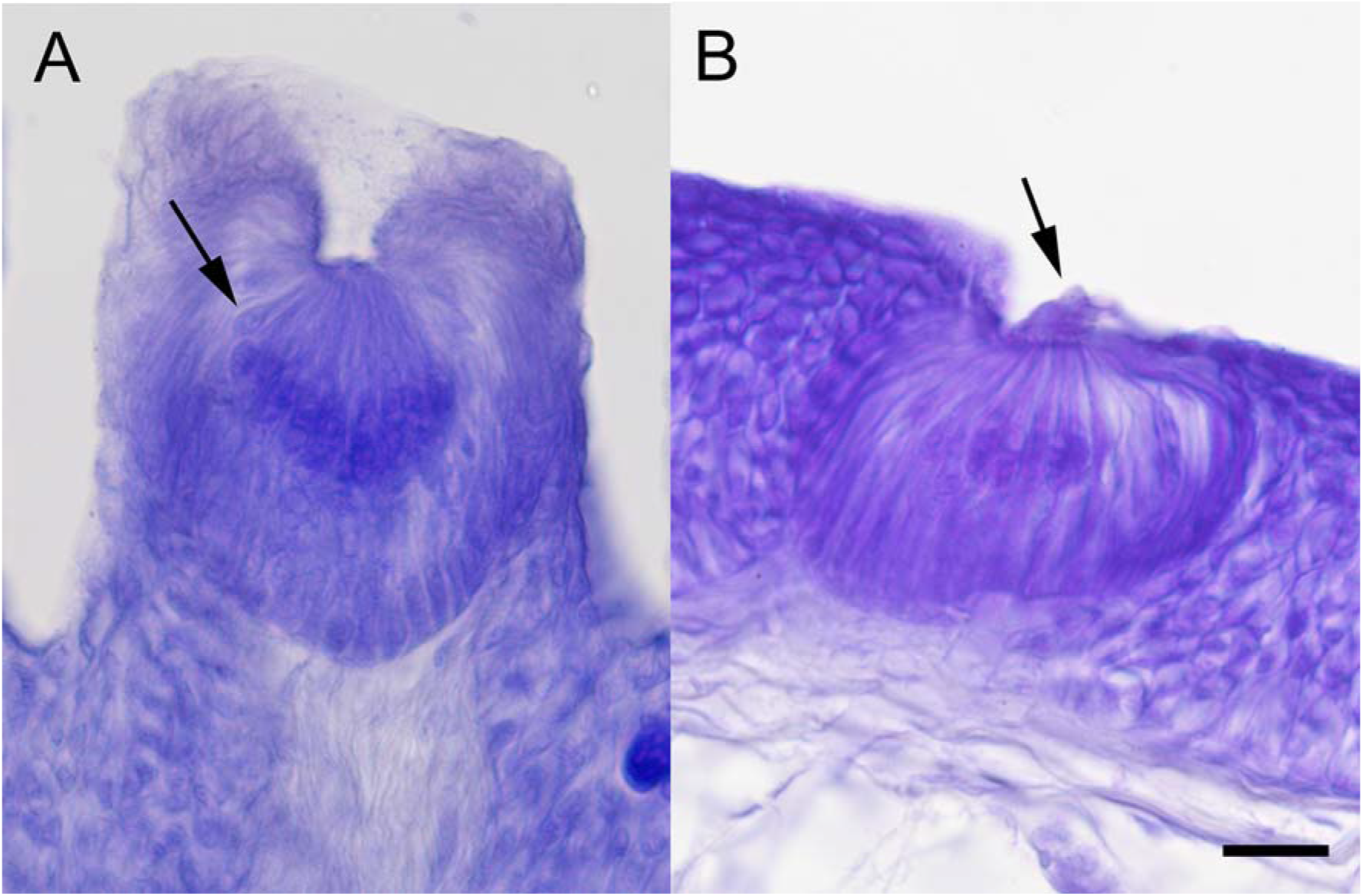
Cross-sections of both types of neuromasts. A) Papillate neuromast from *Amblyopsis spelaea*, arrow points at a single neuromast cell body. B) receded neuromast from *Chologaster cornuta*, arrow points at the collapsed cupula. Scale bar 100um.

*Amblyopsis spelaea* had significantly larger neuromast areas than all other amblyopsids (HSD: P<0.01; Figure 8). There was no difference among *T. subterraneus, F. agassizii and C. cornuta* (ANOVA: P<0.0001, F=70.34, df =28). Sensory plate areas were not significantly different between the two surface fishes, but were both larger than *T. subterraneous* (HSD: P<0.01), and smaller than *A. spelaea* (HSD: P<0.01). Accordingly, *A. spelaea* had a larger sensory plate area than *T. subterraneus* (HSD: P<0.01: ANOVA: P<0.0001, F=62.53, df=28)*. A. spelaea* had longer kinocilium than all other fishes (HSD: P<0.01). *Typhlichthys subterraneus* had longer kinocilium than both *F. agassizii and C. cornuta* (HSD: P<0.01), which were similar in length (ANOVA: P<0.0001, F= 108.95, df= 131). Hair cells in the sensory plate of cavefishes also appear thicker than surface fishes.

**Figure 8.**
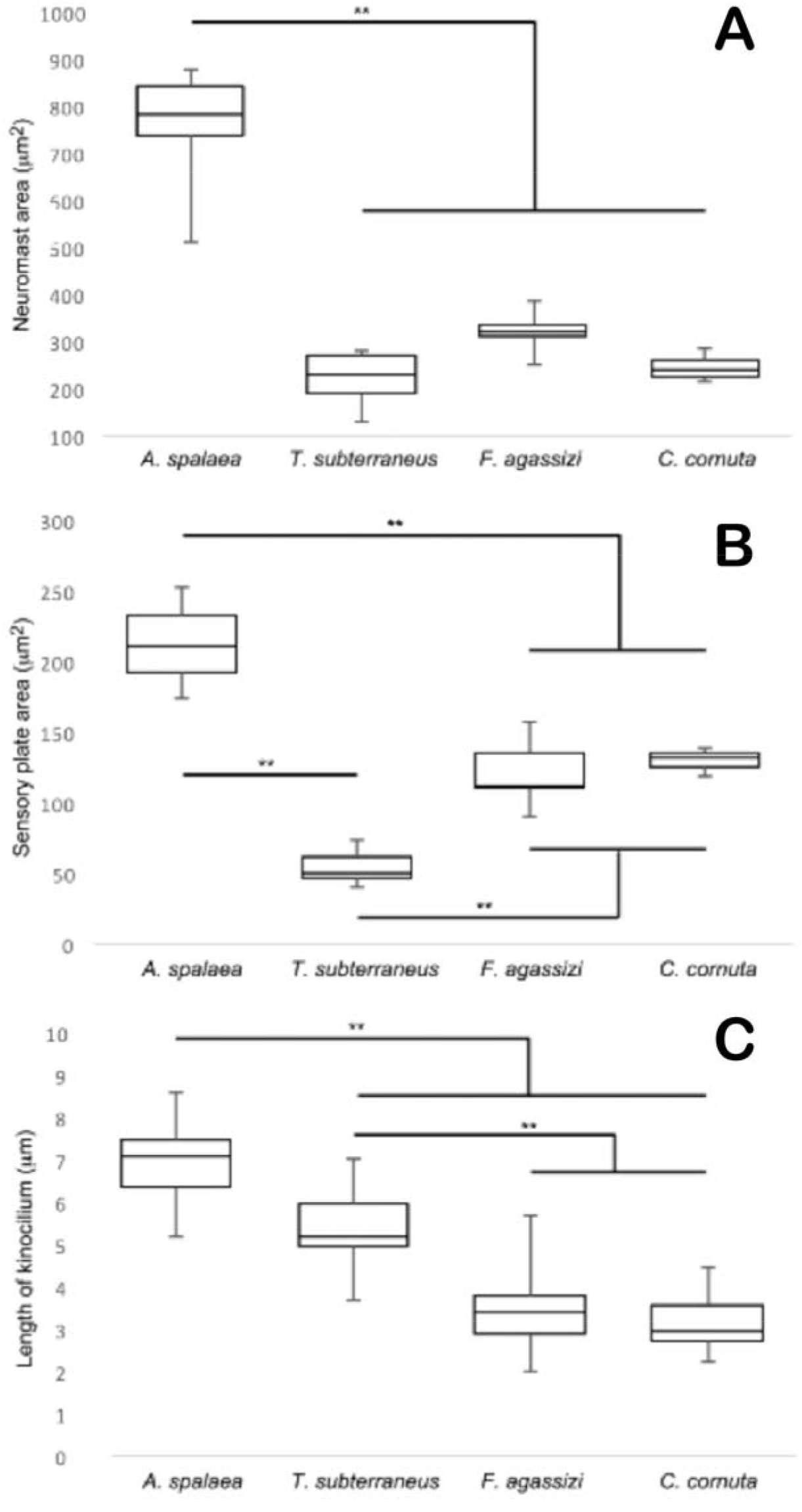
*A. spelaea* has the largest superficial cephalic neuromasts of all fishes (A), with the largest sensory plate area (B) and the longest kinocilium (C). Both surface fishes sensory plate areas (B) and kinocelium lengths are similar to each other (C). Both cavefishes are not significantly different from each other but different from both cavefishes.

## Discussion

The successful colonization of the subterranean environment often is accompanied by the loss of vision. As eyes become less useful for detecting the environment, other sensory modalities such as mechanosensation become more prominent and even hypertrophied (Marshall 1965, 1971). All four amblyopsid fishes included in this study had extensive cephalic superficial lateral line systems. Two types of cephalic superficial neuromasts were present in amblyopsids: one receded into pits into the skin and one on top of papillae. This difference perhaps represents different biological strategies or the result of different environmental adaptations. Both types of superficial neuromasts are arranged in dorsal-ventral stiches on the head. As most of these sets of neuromasts are also arranged with best sensitivity perpendicular to the row, they give the whole lateral line a bias for detecting water movements in the rostral-caudal direction. There were no canal neuromasts seen on the heads of all amblyopsids. In general, proliferation of superficial neuromasts in surface-dwelling fishes is accompanied by the loss of canals (More and Burris 1956; Webb 1989).

The difference is position of cephalic superficial neuromasts between primarily surface and cave-obligate amblyopsids, i.e., receded in pits versus positioned on top of papillae, may reflect differences in microhabitat use. Both *Chologaster* and *Forbesichthys* are nocturnal benthic predators that seek shelter during the daytime and exhibit strong thigmotaxis (Weise 1957; Smith and Welch 1958; Niemiller and Poulson 2010). *Chologaster* typically occurs in heavily vegetated and shaded swamps and backwater habitats where they can be collected in dense vegetation, leaf litter, and other organic debris (Cooper and Rohde, 1980; Ross and Rohde, 2003) and are rarely found in flowing water (Poulson 1960). Likewise, *Forbesichthys* typically are found in dense vegetation, under rocks, or seek shelter underground deep in springs and caves (Weise 1957; Smith and Welch 1958; Niemiller and Poulson 2010). The recession of superficial neuromasts into pits may be an adaptation for living among dense vegetation, organic debris, and other cover in these two species, which could be damaged if exposed and extended from the body surface on papillae.

The difference in position of the neuromasts in the skin in relation to the water is likely to have a functional significance. The water flow that is detected by the superficial neuromasts is filtered by the hydrodynamics of the body surface of the fish (McHenry et al. 2008). The viscosity of water causes flow close to surface to move slower creating a spatial gradient known as the boundary layer (Lamb 1945; Schlichting 1979). The boundary layer over the surface of the body of the fish behaves as a high-pass filter that attenuates low-frequency stimuli (Kuiper 1967; Hassan 1985; Kalmijn 1988; Teyke 1988; McHenry et al. 2008). High frequency stimuli, such as that created by swimming prey or conspecifics, would become more salient to the fish. The boundary layer over the surface of the body of a fish plays a major role in determining the signals detected by superficial neuromasts (Batchelor 1967; Schlichting 1979; McHenry et al. 2008).

The boundary layer of swimming fish was studied in *Astyanax*, and it decreases with velocity so that, it is 5 mm thick at 11 cm/sec, and 1 mm at 150 cm/sec (Teyke 1988). *Astyanax* is comparable in body size to amblyopsids, and, although there are no measurements of swimming speed of amblyopsid cavefishes, the height of the papilla (*A. spelaea* = 0.55±0.006 mm; *T. subterraneus* 0.73±0.16 mm) suggests that the neuromasts likely do not escape the boundary layer. The attraction to live prey using the lateral line has been examined by Yoshizawa et al (2010) also in *Astyanax*. In these fish, the vibration attraction behavior is tuned to 35Hz, which is considered a high frequency and could denote a moving prey item. This behavior is abolished if the superficial neuromasts are ablated, specifically in the cephalic area.

The proliferation of superficial neuromasts is an adaptation for life not only in subterranean environments, but also in the deep sea, and other low-flow habitats (Denton and Grey 1988, 1989; Coombs et al. 1988, 1992; Northcutt 1989; Marshall 1996; Poulson 2001). Deep-sea fishes are an interesting case because their visual world is very different than that of cavefishes. At depth, bioluminescence point sources dominate as visual stimuli, and eyes, although often referred to as ‘regressed’ or ‘degenerate’ are actually quite well suited at this type of stimulus (Douglas et al. 1998; Warrant 2000; Warrant and Locket 2004). Nonetheless, it appears that deep-sea fishes rely heavily on their lateral line to detect the environment and prey, (Marshall 1996, Johnson et al. 2009). Both cavefishes and deep-sea fishes do not display schooling behaviors, which require lateral line input (Cahn et al. 1968; Greenwood et al. 2013; Kowalko et al. 2013). Another group with significant cephalic surface neuromasts are gobies (Marshall 1986; Webb 1989), and in these fishes, many of the stiches are also vertically oriented (More and Burris 1956), biasing the detection of environmental disturbances in the rostral-caudal dimension.

Although we were not able to measure the cupula of the surface species, both cavefishes had a relatively short cupulae, roughly 30 μm in length. The range of lengths of the cupula in surface fishes is variable dependent on the species and ranges between 20 μm and 500 μm, (Mukai and Kobayashi 1993). The cupulae of *Astyanax* are primarily 100 μm in length but some are as long as 300 μm (Teyke 1990). The longer cupulae in *Astyanax* therefore can increase lateral line sensitivity by protruding into the boundary layer. *Astyanax* and amblyopsid cavefishes appear to have different strategies to increase the height in which the cupula interacts with the water. *Chologaster* and *Forbesichthys* were not different from each other, but cavefishes (*A. spelaea* and *T. subterraneus*) appear to encode hydrodynamic information to a different extent. *Amblyopsis spelaea* has the most extreme cephalic superficial neuromast phenotype.

Our study support the general rule that mechanosensation becomes hypertrophied during cave adaptation of fishes. However, our study also suggests that hypertrophy can be variable and possibly dependent on microhabitat, developmental constraints and/or phylogeny.

